# Cytomulate: accurate and efficient simulation of CyTOF data

**DOI:** 10.1101/2022.06.14.496200

**Authors:** Yuqiu Yang, Kaiwen Wang, Zeyu Lu, Tao Wang, Xinlei Wang

## Abstract

Recently, many analysis tools have been devised to offer insights into data generated via Cytometry by time-of-flight (CyTOF). However, objective evaluations of these methods remain absent as most evaluations are conducted against real data where the ground truth is generally unknown. In this paper, we develop Cytomulate, a reproducible and accurate simulation algorithm of CyTOF data, which could serve as a foundation for future method development and evaluation. We demonstrate that Cytomulate can capture various characteristics of CyTOF data and is superior in learning overall data distributions than single-cell RNA-seq-oriented methods such as scDesign2, Splatter and generative models like LAMBDA.

## Background

Recent years have seen exponential growth in applications of mass cytometry (CyTOF) which characterizes proteomics profiles of tens of thousands or millions of heterogeneous single cells on 30-40 protein channels with minimal spill-over effects (1–3). As the technique of CyTOF matures, many data analysis methods have been proposed to offer insights into various aspects of single-cell experiments (4). For example, in (5), the authors have observed that the signal intensity of a CyTOF experiment would vary over time due to changes in instrument performance. Their proposed method is currently the dominant tool to alleviate the temporal effect using polystyrene beads embedded with metal lanthanides. On the other hand, methods such as CytofRUV (6), and Cytonorm (7), etc. aim at getting rid of batch effects to facilitate comparisons across different CyTOF experiments. Deep learning algorithms such as residual networks (ResNet) (8) and generative adversarial networks (GAN) (9,10) were also adopted to resolve the same problem. While some software such as FlowSOM (11), LAMBDA (12), and Bayesian trees (13) focus on automatic cell type identification, others like CytoTree (14) attempt to explore the trajectories along which cells differentiate into other cells. Various data visualization and dimension reduction algorithms have also been applied to CyTOF (15). Typical examples include: t-SNE (16), UMAP (17), MDS (18), SAUCIE (19), scvis (20), etc. As the output of a CyTOF acquisition software is in Flow Cytometry Standard (FCS) format, packages such as flowCore (21) and Seurat (22) have incorporated standardized and efficient pipelines to read, write, and preprocess CyTOF data.

Despite the vast array of methods applicable to CyTOF, most of the CyTOF analysis methods developed by researchers are often evaluated against real data. Without a gold standard and known truth, such as labels and actual distributions, performance assessment becomes somewhat subjective while also blurring the line between method validation and result discovery. The few studies with simulated datasets are highly specialized and workflow-dependent. For example, CytoGLMM (23) focuses on a single cell type for the purpose of differential expression analysis. In (24), the authors further used the simulated datasets in (23) to evaluate performance on various differential expression detection methods. (25) proposed a Bayesian hierarchical model based on Gaussian distributions to study the difference in the protein functional network of a single cell type under two experimental conditions. In (26) the simulation was done by manually splitting a real dataset into two overlapping subsets. These simulation procedures are not specifically designed for general-purpose simulations of CyTOF samples with many cell types, trajectories, batches, etc.

Simulated datasets, albeit being synthetic, contain ground truth against which methods and algorithms can be reliably tested. Instead of *ad hoc* simulation in each study mentioned above, a formal simulation tool naturally provides a framework for a systematic evaluation for new methodological research in the field. By varying parameters and settings of key factors, researchers can easily generate an unbiased coverage of the system under study in a straightforward manner. On the other hand, real experiments require detailed prior planning, precise executions, and potentially abundant resources. The flexibility along with cost and time benefits make simulation a vital tool for the field. However, the lack of a rigorous, reliable simulation algorithm calls for the development of such a tool for validating other CyTOF data analysis tools and future developments.

By contrast, various methods have been developed for simulating single cell RNA-sequencing (scRNA-seq) data, such as Simple (27), Lun (28), BASICS (29), scDesign2 (30), ZINB-WaVE (31), and Splatter (32), among which Splatter enjoys the highest popularity whilst scDesign2 excels in simulation performance (33,34). Since both scRNA-seq and CyTOF are single-cell sequencing techniques, it might be tempting to tweak scRNA-seq simulation tools such as Splatter and scDesign2 and apply them to CyTOF. However, such a process can be highly nontrivial and the end results may be unsatisfactory if only small adjustments are sought after. First of all, these two techniques are of different biological nature. scRNA-seq examines the expression of thousands of RNAs of several thousand cells, whereas CyTOF captures information on typically 30-40 proteins for tens of thousands or even millions of cells. Therefore, compared with scRNA-seq, CyTOF is more closely related to clinical phenotypes and is more likely to identify rare cell populations. Furthermore, while drop-out is a common issue in scRNA-seq data, it is a much less encountered problem in CyTOF experiments. The expressions in scRNA-seq are integer-valued “counts”, in contrast to real-valued data that even have negative values in CyTOF expression matrices due to several built-in processing steps in a CyTOF system such as data randomization and transformation. Consequently, applying simulation tools like Splatter and scDesign2 will almost inevitably require a sequence of nonlinear transformations to artificially construct count data, which will yield biased results. Finally, the expression levels of various protein channels in CyTOF are naturally dependent. Extending a complex discrete random variable model such as the Poisson-Gamma mixture model adopted by Splatter to capture the dependency structures present in CyTOF may not be a trivial task. In addition, methods for simulating multivariate discrete data such as the ones devised in (35,36) are not suitable for CyTOF data that are real-valued. The same logic applies to other generic simulation algorithms that are not designed with CyTOF in mind.

To bridge this gap in literature, we developed Cytomulate, the first formal simulation tool that is capable of generating simulated datasets that capture various characteristics of CyTOF data. Our framework combines the flexibility of generating complex, realistic data with efficiency and usability, allowing researchers to acquire data at scale and practitioners to easily simulate datasets for their workflows.

## Results

### Characteristics of CyTOF data

Currently, most CyTOF data are available in the FCS format for analysis purposes, containing real-valued protein expression levels rather than counts. This is due to several built-in data processing steps in a CyTOF system (e.g., Helios, latest CyTOF XT): (i) The system applies regression to recover peak counts lost because of overlapping peaks at high intensities, resulting in non-integer count estimates. (ii) To avoid binning effects caused by integers near zero, the system applies randomization (i.e., adding uniform or Gaussian noise to counts). (iii) Bead normalization is applied to remove temporal effects due to changes in instrument performance over time. (iv) The inverse hyperbolic sine function, referred to as arcsinh-transformation, is applied to alleviate the coupling of mean and variance and “gaussianize” the data (37). Although the system allows users to turn on and off each of (ii)-(iv), often one or more such options were implemented for nearly all existing CyTOF data, leading to real-valued expressions.

To motivate the design choices of Cytomulate, we explored prominent characteristics of CyTOF by analyzing various public and in-house datasets (e.g., Finck (5), Levine_13dim (38), Levine_32dim (38), CytoNorm (7), CyAnno (39), Covid (15); Table S1). In this section, we mainly use protein expressions from the Levine_13dim dataset as an example to demonstrate such features. On a broader scale, similar phenomena have been observed across numerous CyTOF datasets. The Levine_13dim dataset is publicly available via the R package HDCytoData (40). As the protein expressions have already been pre-processed to correct unwanted artifacts, such as temporal effects, only arcsinh-transformation was applied.

### High volume, high dimension, and heterogeneity

The expression matrix associated with a typical CyTOF experiment usually contains tens of thousands or even millions of cell events (rows), 30-40 protein channels (columns), from a pool of cells of 10-30 cell types. For example, the data of the first patient in Levine_13dim has 167,044 cell events, 13 protein channels, and 14 cell types manually gated by the authors. With a varying number of protein channels, the heterogeneous nature of different cells are captured by CyTOF via cell types and differentiation stages. Any potential simulation method developed for CyTOF needs to not only handle the throughput with adequate efficiency but also the heterogeneity of cell events.

#### Zero-inflation

The protein expression matrix observed in a CyTOF experiment often contains features with either zeros or values close to zero. These features biologically correspond to surface proteins that are too scarce to be detected by the machine. In addition, because of data randomization, one of the standard options in the CyTOF system, many datasets have near-zero or even negative values rather than zeros. Take, for example the CD4 T cells in the dataset, we see in Fig. 1a and Fig. 1b that the majority of the protein channels (8 out of 13) contain a high proportion of values close to zero.

**Fig. 1.**
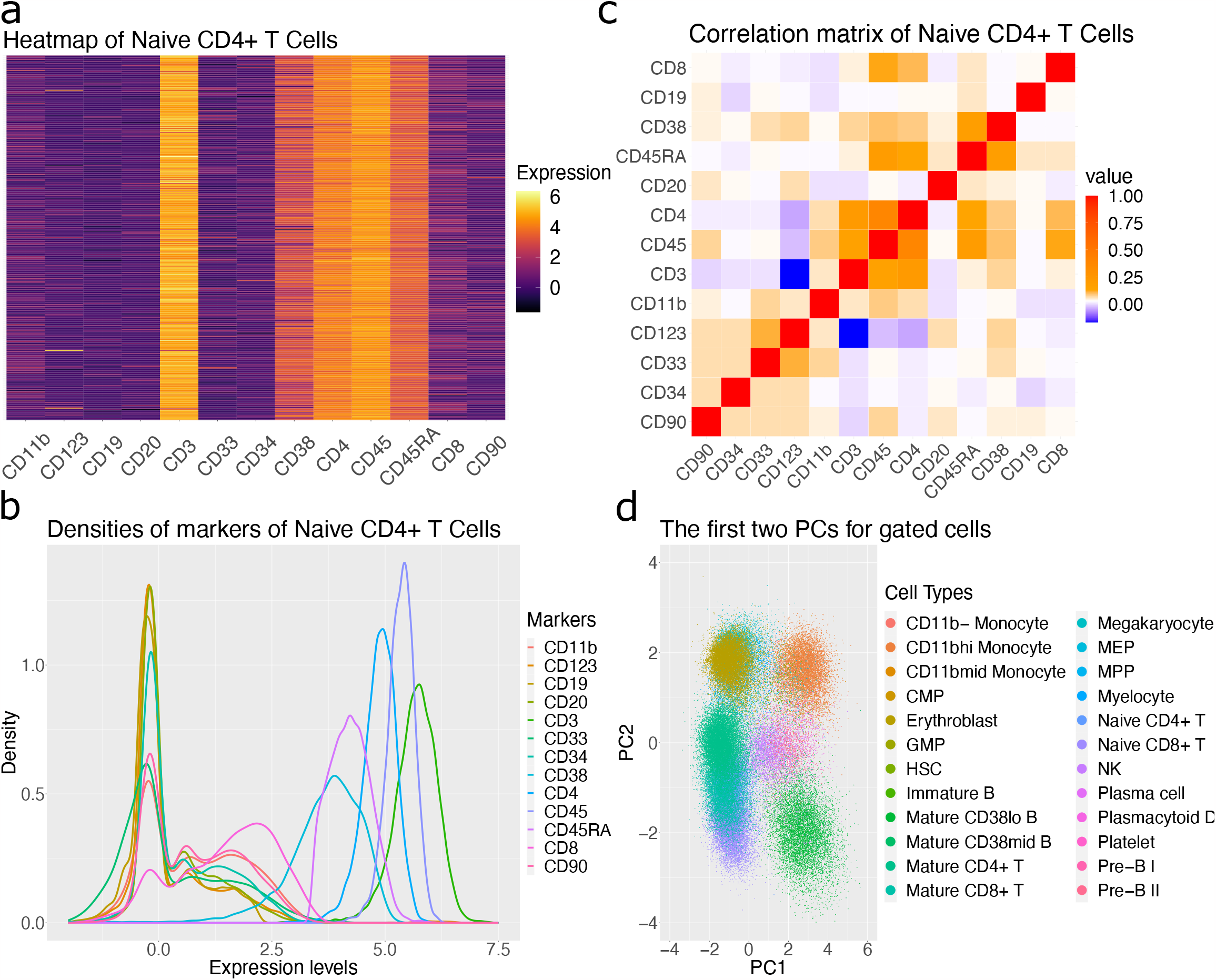
Characteristics of a typical CyTOF dataset. All four panels are from the Levine_13dim as an example. (a) Heatmap of channel expression for all naive CD4+ T cells. (b) Density plot of all channel expressions from naive CD4+ T cells. (c) Correlation matrix of all channels as computed with Pearson correlation coefficients. (d) Scatter plot of all cells using the first two principal components and colored with provided cell types.

#### Correlation in the protein-protein expression

In Fig. 1c, we calculated the correlation matrix among all the protein channels using the data of Naive CD4+ T cells. Clearly, markers such as CD44 and CD45 are correlated, implying that the independence assumption between columns might be too restrictive. This suggests modeling the data jointly instead of marginally.

#### Near-normality

To explore the distributional characteristics of arcsinh-transformed protein expressions, we first examined the marginal density estimates of each protein marker for different cell types. In Fig. 1b, we plotted these empirical densities for all markers using Naive CD4+ T cells, where expressed markers (i.e. those with relatively high expression levels) possess “bell”-shaped curves and other markers exhibit two or more modes that can be approximated by Gaussian mixture models (GMMs). To further get a sense of the joint distribution, we performed dimension reduction using Principal Component Analysis (PCA). All cell populations resemble roughly elliptic shapes (Fig. 1d), suggesting that there is no gross violation to the joint normality, according to the assumptions of PCA. It may be tempting to use a multivariate normal (MVN) distribution for each cell type. However, protein markers’ distributions depend on user-provided cell types and so MVNs may not be adequate, particularly in cases where the cell typing is coarse or inaccurate. Given that GMMs are known as a universal approximator of densities (41), combined with the near-normality of CyTOF data described above, it is a sensible choice to use GMMs with Gaussian distributions as basic building blocks of a simulation model. Finally, this key characteristic also differentiates CyTOF from scRNA-seq, which is typically modeled with counts. As explored in the comparison study later in this paper, adapting a model for discrete counts for CyTOF does not work well.

#### Temporal effect

As illustrated in (5), the signals of a single CyTOF experiment would typically vary over time due to changes in instrument performance. Since Cytomulate aims at modeling arcsinh-transformed protein expressions, unlike in (5), we chose to demonstrate the signal drift by transforming the Peripheral Blood Mononuclear Cell (PBMC) data from the Finck dataset (Fig. S1a). As a first attempt to model the signal drift, we centered each signal, fitted a smoothing spline against one of them, and subtracted the fitted value from each signal. In Fig. S1b, we see that the adjusted signals overlap and are roughly homoscedastic, suggesting that the overall shape of the signal drift (temporal effect) may be shared by all cell types within a single batch and that we could model the effect using an additive model.

#### Batch effect

Apart from the temporal effect, many researchers also reported the presence of batch effects across multiple CyTOF experiments due to sample collection, stimulation, staining, etc. (42). In (7) they reported that not only are the batch effects cell-type specific, they also vary depending on protein channels. In Fig. S1c, we illustrated this phenomenon by an interaction plot using the mean expressions of the protein channels of CD4 T cells and CD8 T cells measured on two patients in the Levine_32dim dataset (40). We see that not only do expression levels differ between two batches, the differences also vary by cell types and protein channels. This implies that in addition to the main batch effect, we should also take into consideration the interaction effects.

#### Cellular trajectory

A common task is to study cellular trajectory where the expressions of an individual cell are assumed to vary according to some continuous path that connects one cell type to another. In the context of scRNA-seq, lots of methods have been developed to align cells on continuous paths with typical examples Slingshot and Monocle (43–45). To our knowledge, CytoTree (14) is the only available method of trajectory inference designed for CyTOF. It is further questionable whether trajectory inference methods are interchangeable between these two technologies. For the purpose of demonstration, we showed an example of the cellular trajectories following CytoTree’s online tutorial with a real Bone Marrow dataset in Fig. S1d.

### Cytomulate framework

In light of the previous exploratory data analysis, we propose Cytomulate, an accurate and efficient simulation algorithm of CyTOF data based on Gaussian Mixture Models (GMM) (46), with add-on features to mimic various patterns/effects existing in real data. More specifically, Cytomulate achieves the task of accurate simulation via two modes selectable by users: emulation mode and creation mode, both offering transparency with known ground truth. Emulation mode is designed for a user to simulate data that aims to capture major characteristics of a specific CyTOF dataset under study. As this mode learns meaningful parameters from real data, it enables interpretation of real data such as locations, variations of protein expressions, protein-protein correlations, and heterogeneity among various cell types, while generating synthetic data to test different hypotheses. Creation mode is purely model-based in which synthetic data are generated from user-specified settings and parameters. This mode caters to two main user groups. Firstly, it serves users who wish to have access to a wide range of CyTOF datasets, encompassing varying numbers of cells, protein markers, experimental designs, tissue types, diseases, and conditions. These users aim to simulate multiple datasets with diverse characteristics to facilitate comprehensive testing and benchmarking. Secondly, creation mode is also designed to assist new researchers in the field who may not have a real CyTOF dataset at hand. By utilizing this mode, they can generate synthetic data immediately, providing an accessible starting point for their own experiments and studies.

In both simulation modes, users are allowed to vary cell abundances, noise levels, batch effects, temporal effects, as well as cellular trajectories to capture the complex needs of methodological developments for CyTOF data. The simulated batch effects, temporal effects, cellular trajectories, as well as the noise are then added once we sample from the GMMs. The following sections elaborate on each mode with greater details as the foundation to the comparison study that follows.

**Creation mode,** aims at quickly generating synthetic datasets that capture aforementioned characteristics of CyTOF data such as zero-inflation, correlation in protein-protein expressions, and near-normality. Users specify the volume, dimension, and the degrees of heterogeneity. Temporal effect, cellular trajectory, and batch effect can be added as part of complex simulations. As this mode does not intend to extract the data distribution of some specific dataset as a reference, it is particularly useful for testing analysis methods on a large scale. For example, in (15), Wang et al. used this mode of Cytomulate to evaluate the performance of 20 dimension reduction methods from various aspects such as local and global structure preservation on 425 simulated datasets with different attributes (e.g., different numbers of cell types and numbers of differentiation paths). This mode allows practitioners to precisely define each simulated dataset as they wish, which is not often or at all possible with real data.

Specifically, the creation mode randomly generates average cell expressions and then utilizes GMMs from which to sample cell expressions. Users also have the options to enable complex simulations by simply specifying the necessary parameters, such as the number of trees for cell differentiation trajectories. The details of complex simulation will be discussed later. With users having full control of their simulated dataset, the creation mode is untethered by existing—yet oftentimes implicit—conditions of real datasets, such as the exact cell types, circumstances of the patient, the physical machine, etc. By eliminating these factors, Cytomulate allows researchers to develop and test methods based on unbiased results that are generalizable to wide ranging real datasets. Further, it can also serve as a pedagogical tool for researchers to explore the characteristics of CyTOF datasets easily.

**Emulation mode**, on the other hand, is designed to learn the distribution of some real CyTOF data. To ensure compatibility, Cytomulate assumes that the reference dataset provided by the user has already undergone bead normalization and arcsinh-transformation (with a cofactor of 5), both of which are standard options in the CyTOF system. Instead of approximating the entire dataset as a whole, which is challenging, Cytomulate leverages the prior knowledge of cell type information. As a result, it requires not only the expression matrix of one particular CyTOF dataset to mimic but also the cell type label associated with each cell as inputs. Although Cytomulate can potentially perform cell type identification or clustering using common algorithms of the likes of K-means, we leave the choice of such algorithm for cell typing to our users as a design choice for the following two reasons: (i) According to (47) and (48), the k-means algorithm and GMM-based model (SWIFT) are often not the optimal choice for clustering CyTOF data; (ii) There are already plenty of well-established cell identification algorithms and automatic gating techniques such as FlowSOM (11), Bayesian Tree (13), ACDC (49), SCINA (50), etc., all of which have similar goals but slightly different design considerations. For example, using FlowSOM requires minimal prior knowledge on the cell population other than some crude idea on the number of cell types. On the other hand, Bayesian Tree and ACDC use a template as an input to facilitate the automatic gating process. Since we expect the users of the emulation mode to be somewhat experienced in the field, we think it would be better to let the users choose and apply their own cell identification algorithms that suit their situations the best. The resulting expression matrices of this mode are thus expected to be a closer approximation to the real data, as opposed to those generated by the creation mode.

To dive into the details, Cytomulate, at its core, uses a GMM with truncation below 0 to approximate and generate the probability density of a given cell type. The Bayesian Information Criterion (BIC) (51) is adopted for model selection. Although the exploratory data analysis in the previous section shows that a MVN might be a good candidate for modeling a specific cell type in a CyTOF experiment, this should be taken with a grain of salt because the cell-typing procedure can vary across datasets and cohorts. For example, the Levine_32dim dataset provides labels for the following subtypes of B cells: mature B cells, plasma B cells, pre-B cells, and pro-B cells. In practice, however, subsets of major cell types are not available in certain datasets. We thus should allow flexibility in the model to account for cell-type procedures with different resolutions. If the users are interested in subtypes of B cells, a single MVN would be inadequate in describing the data generating mechanism (see the comparison study in the next section for details). Therefore, we chose GMMs as the base for its flexibility and its mathematical tractability.

In Fig. 2a, we outlined the overall framework of Cytomulate with input and output examples, model specification, and possible workflows for downstream analyses. Further details on the two simulation modes and the probabilistic model can be found in the Materials and methods section.

**Fig. 2.**
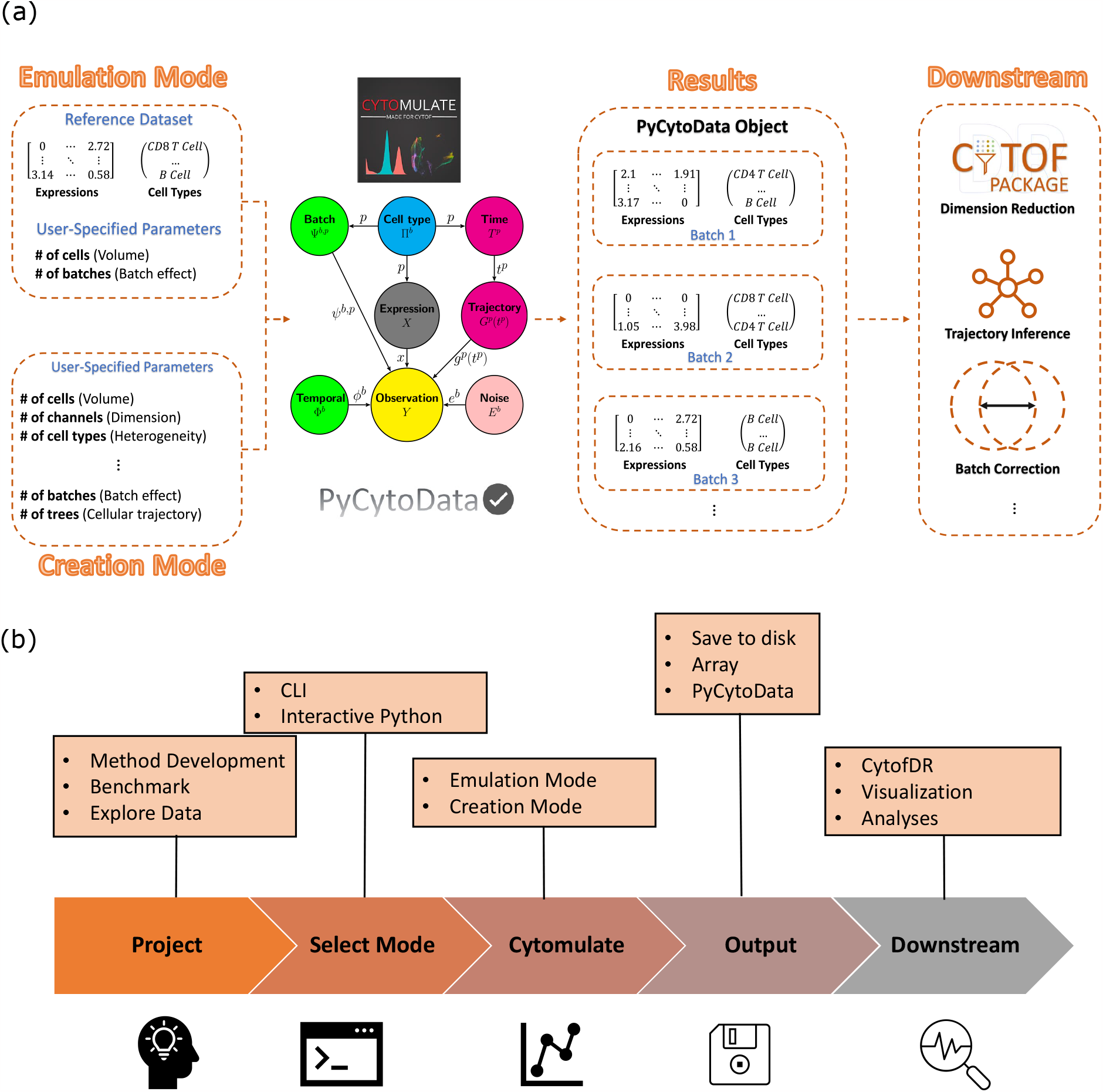
Overview of Cytomulate model and software pipeline. (a) The model structure and input and output pipelines of Cytomulate. The input stage demonstrates the parameters and reference data needed for each mode, and Cytomulate uses the provided information accordingly for estimation and simulation. The outputs are in the forms of expression matrices and their associated cell types for additional downstream analyses. (b) The software pipeline of Cytomulate. The flowchart showcases Cytomulate’s role and integration with industry standard tools in every step of the research and development process.

We also modularized all the aforementioned functionalities so that each of them can be easily fine-tuned or turned off with a simple interface, according to a user’s specific need. Cytomulate returns the final expression matrices, the cell type labels associated with the expression matrices, as well as cellular trajectory information for each cell event. Optionally, users can also choose PyCytoData as the output option for downstream analyses compatibility (Fig. 2b). In the following sections, we conduct some comparison studies and demonstrate the usage of Cytomulate through examples.

### Simulation comparison study

Although Cytomulate is the first-of-its-kind simulation tool designed specifically for CyTOF experiments, we will be remiss not to include benchmarks since it’s a common practice in the field to adopt similar technologies from other fields to CyTOF. To showcase the superior statistical and computational properties of Cytomulate, we performed a series of empirical experiments using the emulation mode. In this simulation setting, the real benchmark datasets provide a gold standard for assessing the accuracy of each simulation method and the overall validity of the underlying model. More specifically, we focused on whether each model is capable of capturing the features of CyTOF datasets and the overall distributions and relationships between all protein channels.

For the comparison study, we included the following models from within Cytomulate and other fields:

1. Cytomulate with full covariance and model selection via Bayesian Information Criterion (BIC) from GMMs with components varying from 1 to 9 (Cytomulate)
2. Cytomulate constrained to one Gaussian component with full covariance per cell type (Dependent Gaussian or simply DG)
3. Cytomulate constrained to one Gaussian component with diagonal covariance per cell type (Independent Gaussian or simply IG)
4. Splatter (32)
5. Latent Allocation Model with Bayesian Data Analysis (LAMBDA) (12)
6. scDesign2 (30)

As discussed in the **Characteristics of CyTOF data** section, the correlation of protein-protein expressions is an important feature of CyTOF datasets. We thus included two reduced versions of Cytomulate, namely IG and DG, both featuring only one multivariate normal component per cell type. The IG model focuses on the diagonal covariance matrix to investigate the effect of omitting the covariance structure seen in Fig. 1c. The DG model serves as a direct comparison against IG by considering the linear dependencies among protein channels. Finally, our core model, Cytomulate, aims to showcase the advantages of modeling non-normality through multiple normal components per cell type, in contrast to DG. The inclusion of these progressively complex models will not only demonstrate the flexibility of the Cytomulate framework, but it will also validate the characteristics of CyTOF, as shown in the exploratory data analysis, by using actual benchmarks.

Beyond Cytomulate and its variants, we also included Splatter, LAMBDA, and scDesign2, which are not designed for CyTOF simulation. Below we give a brief description of each followed by the rationale for their inclusion in our comparison study.

**Splatter**, or specifically Splat, is originally designed for simulating scRNA-seq data. Its core model uses a Gamma-Poisson mixture to simulate counts observed in scRNA-seq experiments. The authors also take into consideration outliers, library size, mean-variance trend, and technical dropout. As Splatter expects integers as inputs, to simulate CyTOF data which usually have been transformed into a numeric scale using the arcsinh transformation, we can carry out its inverse transformation, round up the results to the nearest integer (37), apply Splatter to simulate counts, and transform the simulated expressions back into the numeric scale. We included Splatter to demonstrate the consequences of adapting scRNA-seq simulation methods to CyTOF, which outweigh the tempting benefit of convenience for any rigorous CyTOF studies.

**LAMBDA** is a Bayesian hierarchical model designed for automatic cell identification in CyTOF data. The model assumes that given a cell type, the cell expressions follow a zero-inflated multivariate Gaussian distribution. The cell types, on the other hand, are generated from a Categorical distribution whose parameters are controlled by some clinical information.

Parameter estimation is accomplished by a stochastic EM algorithm. It is worth pointing out that LAMBDA is not a simulation tool and its simulation functionality is only a byproduct of LAMBDA. More critically, LAMBDA cannot generate new datasets without using reference datasets. Thus, for those who are interested in the Creation Mode or those who do not have access to any previous samples (*e.g.* CyTOF for some rare diseases), this is not a suitable method at all. We included the method in our study because of its rigorous parameter estimation procedure, which would provide us with a yardstick against which we can evaluate the quality of the simulated datasets in terms of the first and second moments.

**scDesign2** is an scRNA-seq simulation algorithm designed to address the shortcomings of other methods in the field, including Splatter. This algorithm first fits one of four discrete distributions (zero-inflated negative binomial, negative binomial, zero-inflated Poisson, or Poisson) to each gene’s marginal distribution. Then, it employs a Gaussian copula for each cell type to achieve joint modeling, retaining the gene expressions from the real data while also capturing the correlation structure within each cell type. The key difference between scDesign2 and Cytomulate is that scDesign2 is specifically designed for discrete data, which presents similar challenges to the application of Splatter for CyTOF data.

To demonstrate the versatility of Cytomulate, six publicly available datasets from different species and diverse anatomic sites are collected: Levine_32dim (38), Levine_13dim (38), Samusik (52), CyAnno (39), Covid (15), and LG (53). If multiple samples exist for one dataset, the first sample is used. See Table S1 for details. All of these datasets have been bead-normalized and transformed by using the arcsinh transformation. Cell type information is also publicly available for the first four datasets. Cell types in Covid and LG were identified via FlowSOM and manual gating (15).

### Cytomulate well approximates real data

In this comparison study, we make use of all six datasets and include the aforementioned six models as candidates. Four metrics are adopted to quantify the quality of the simulated datasets: 1. L2 distance between the simulated mean and the observed mean; 2. Frobenius distance between the simulated covariance and observed covariance; 3. Kullback-Leibler (KL) divergence (54) between the simulated and observed channel expressions; 4. Propensity Mean Squared Error (pMSE) (55) between the simulated and observed data. The L2 distance and the Frobenius distance are defined at the cluster level, whereas the KL divergence and pMSE are computed at the single-cell level for each cell type without using summary statistics. For pMSE, we report the relative efficiency (RE) of each competing method over Cytomulate, defined as the ratio of the pMSE value of Cytomulate vs. that of the other method. Here, for any specific method, a RE value larger than one indicates this method is more efficient than Cytomulate in terms of pMSE and by definition, the RE value for Cytomulate is always one. We also ranked the six methods based on raw values of each metric, where the best methods received an order of 6 with the worst being 1. See the Materials and methods section for further details. For each dataset, the process is repeated 50 times for 50 simulated datasets.

Using the Levine_32dim dataset, detailed comparisons show that Cytomulate outperformed all other methods benchmarked in this study in the categories of KL divergence and pMSE (Fig. 3a). As the KL divergence is a measure of similarity between two distributions, Cytomulate’s resounding lead is a testament to its ability to yield a better approximation to the real data compared with other methods. Cytomulate’s advantage in distributional properties logically explains its lead in pMSE, which measures whether simulated datasets are distinct from real ones. Overall, for mean estimation, all methods except Splatter and scDesign2 work well (Fig. 3b-c), with LAMBDA achieving the smallest Mean Square Error (MSE) in some datasets, closely followed by Cytomulate. Even though LAMBDA’s estimation procedure should theoretically yield decent results for the first two moments, Cytomulate still outperformed LAMBDA in estimating the covariance structure in some datasets (Fig. 3b). This result also empirically demonstrated that potential improvements can be made if only the first two moments are considered, but this improvement—if having any practical significance at all—may come at the cost of worse KL Divergence and computational efficiency. Splatter, due to the usage of nonlinear transformations and marginal modeling, performs the worst in all benchmarks by an overwhelmingly wide margin, demonstrating that Splatter is suboptimal when simulating CyTOF data. Although scDesign2 performs joint modeling, which contributes to its superior performance against Splatter, the discrete nature of its model makes it a poor choice overall.

**Fig. 3.**
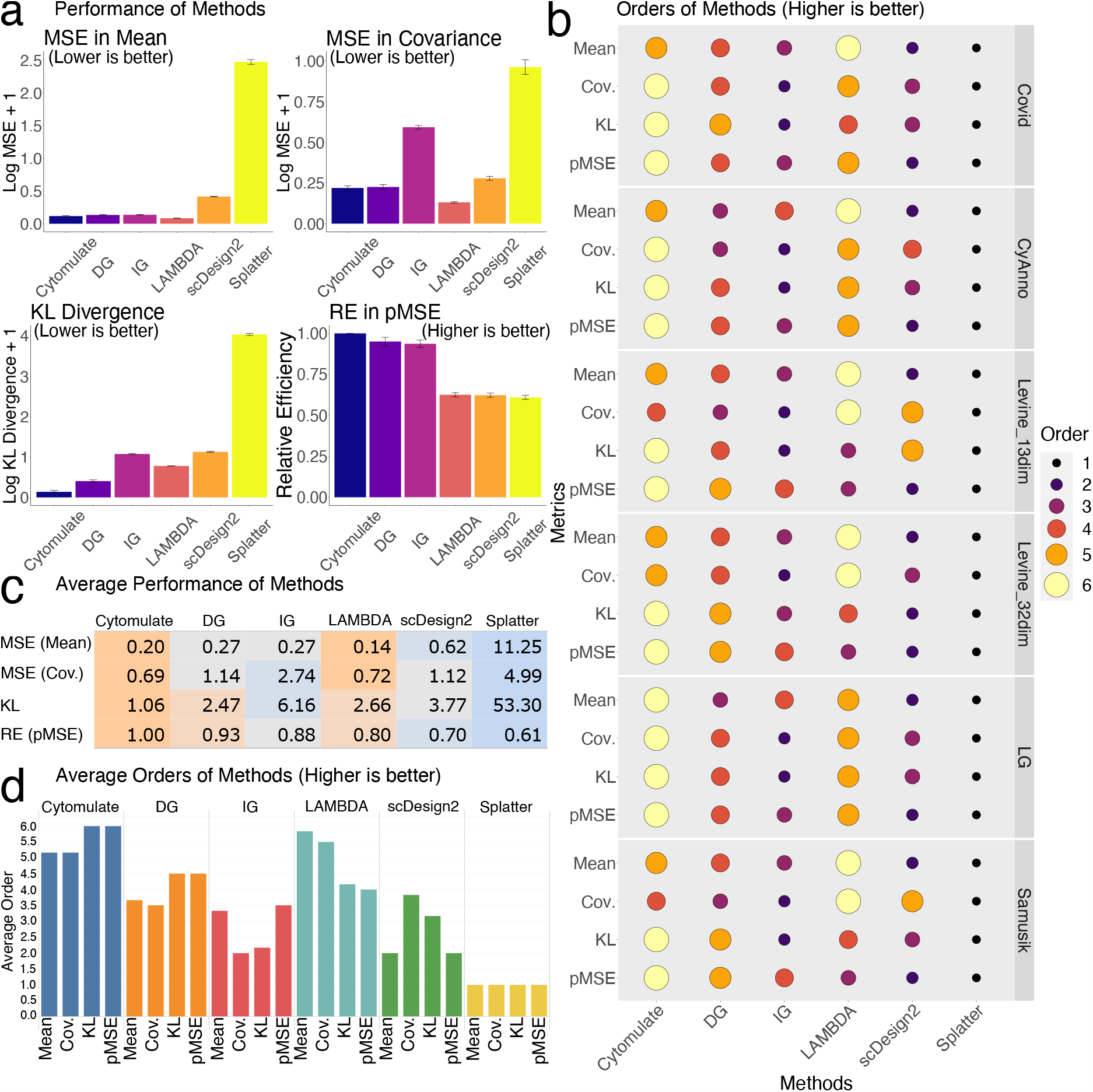
Detailed results on simulation performance benchmarks. (a) The performance of all methods in the Levine_32dim dataset. The four panels correspond to MSE in mean (smaller is better), MSE in covariance (smaller is better), KL Divergence (smaller is better), and relative efficiency in pMSE (larger is better). The relative efficiency is measured using Cytomulate’s pMSE: values smaller than 1 indicate worse performance than Cytomulate. (b) Orders of each method across all datasets and all metrics. Orders are based on values from the metrics, and a higher order indicates superior performance in the given metric. (c) The average performance of methods averaged across all datasets. Orange colors indicate superior performance, whereas blue colors represent worse performance. The interpretations are the same as (a). (d) Orders of all methods averaged across all datasets. Higher bars represent a higher average order for a given metric and method. Colors represent different simulation methods.

Aggregating results from all six datasets using all four metrics, the overall trend observed previously holds for Cytomulate (Fig. 3c). By using GMMs to fit and sample from each cell type, Cytomulate gives the best approximation to the real data as measured by the KL divergence and pMSE. Averaging the orders in Fig. 3b across six datasets further highlights Cytomulate’s advantages (Fig. 3d). As expected, by formally deriving a rigorous estimation procedure, LAMBDA is able to achieve the best average MSE in terms of mean while its covariance estimation lags slightly behind. However, as we have seen previously, the sufficiency of only considering the first two moments highly depends on the data and the coarseness of the cell type labels. It is also evident that Splatter always gives the worst performance due to the bias introduced by nonlinear transformations. Even working with the simple IG would yield a performance boost. scDesign2 outperforms Splatter on all metrics, but it still falls behind Cytomulate variants and LAMBDA.

To visually validate the performance of each method, we applied UMAP (17) to illustrate the resulting expression matrices of each method (Fig. S2). Due to the granularity of the cell type labels included in the dataset, at the first glance, all methods except for Splatter produce reasonable simulations. Splatter produces a concerning embedding with cells splattered around the clusters and cell types indistinguishable. For all other methods, a closer examination reveals that only Cytomulate is able to preserve granular details in the simulation while other methods present only the big clusters. The preservation of detailed characteristics provides an advantage not only in preserving rare cell types and their unique phenotypes, but also in situations where cell-type resolution is low, which motivates our investigation in the following section.

### Cytomulate is robust against cell-type resolution

Although cell type labels are required inputs for the emulation mode, Cytomulate does not demand highly granular labels, meaning that it is not necessary to identify all subtypes of a given cell type as a prerequisite for the simulation. In fact, different datasets and analyses call for different cell typing approaches: a large dataset with only B cells may warrant more detailed cell types to reveal their differentiation stages, whereas small whole blood samples may not afford the same resolution with reasonable cluster sizes. To account for these two situations, we conducted two separate benchmarks regarding cell typing. We first focused on a dataset with many cell types and then zeroed in on B cells to investigate the effects of subtypes.

In real data, the presence of unidentified or unassigned cells due to factors like noise, specific algorithms, and instrument panels is common, making it important to address such scenarios for reasonable results. To simulate this situation, we randomly masked some cell types in the Levine_32dim dataset and assessed the KL divergence between the simulated and original datasets. Specifically, we conducted a benchmark with three settings, randomly selecting and treating 2, 5, or 10 cell types as the same type, reflecting cases in which these cells are not properly identified despite potentially distinct properties. Our results show that Cytomulate is unequivocally the best performer among the six methods tested (Fig. 4). As expected, when cells are unmasked (i.e. cell labels are correctly specified), the KL divergence is smaller, but the procedure of masking cells only yields a small increase in KL divergence for Cytomulate (Fig.4). As the number of masked cell types increases, Cytomulate’s advantage is further accentuated as compared to LAMBDA and scDesign2’s declining performance (Fig. 4b-c). In contrast, Splatter consistently performs poorly, and the issue is further amplified when masking cell types (Fig. 4).

**Fig. 4.**
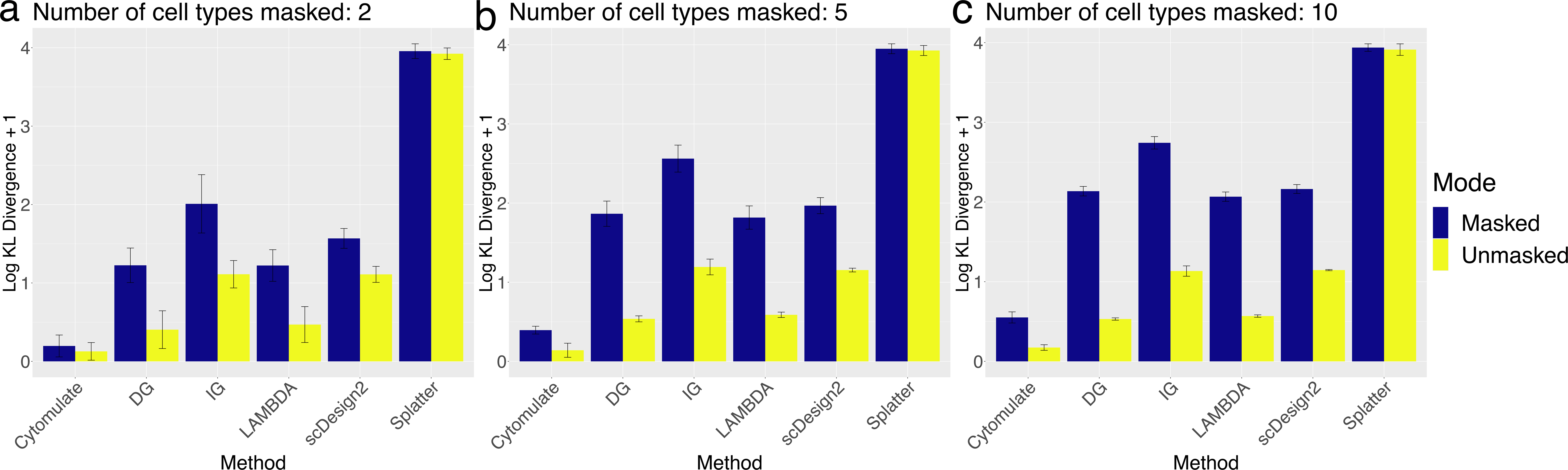
Performance of simulation methods against random masking of cell subtypes. For all three panels, randomly cell types of the Levin_32dim were masked and performance of each method was measured using log KL divergence plus 1. Blue bars represent the masked datasets, and yellow bars represent matched, unmasked controls (N=20). The whisker on top of the bars indicates the variance. (a) 2 cell types are randomly masked in each replicate. (b) 5 cell types are randomly masked in each replicate. (c) 10 cell types are randomly masked in each replicate.

We have shown Cytomulate’s advantage when potentially dissimilar cell types are masked. Now we proceed to investigate the effect of masking similar subtypes. As we mentioned previously, there are 4 subtypes of B cells in Levine_32dim dataset: mature B cells, plasma B cells, pre-B cells, and pro-B cells. In this benchmark, we first provided every method with these detailed subtype labels, and then amalgamated all the labels into “B cells”, masking the subtype details. To quantify the differences observed across the methods, we repeated the process 20 times and computed the KL divergence (54) between the simulated and real datasets under the two settings (Fig. 5). Although the overall performance is consistently improved in the unmasked setting with more detailed cell subtypes, Cytomulate appears to be the most robust among all methods. In fact, both settings of Cytomulate outperformed all other methods regardless of cell-typing details.

**Fig. 5.**
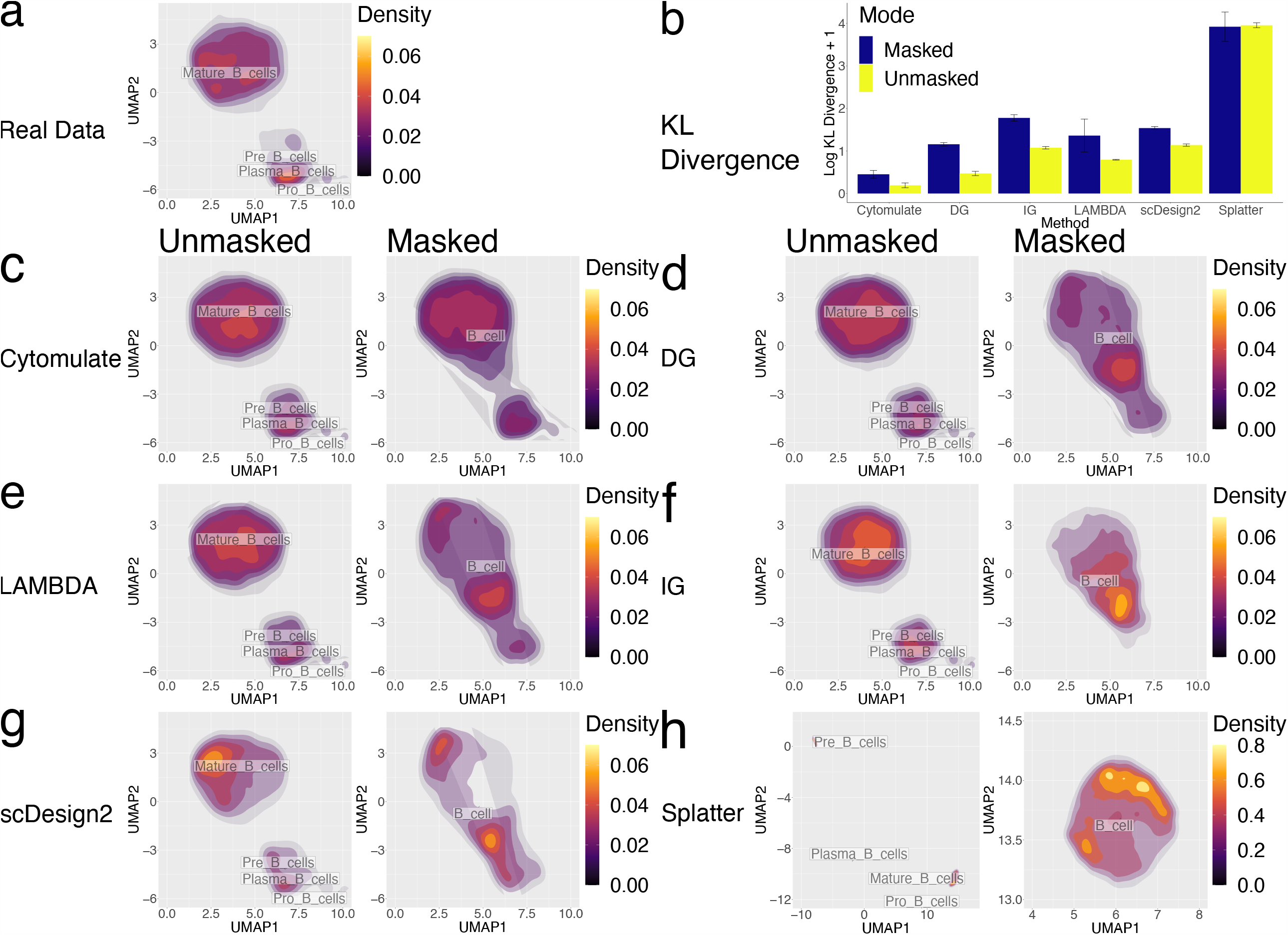
Additional results on performance with different cell subtypes. All parts of the figure utilizes the Levine_32dim dataset with B cells only. (a) UMAP embedding of the original B cells present in the Levine_32dim dataset overlaid with B cell subtype labels as well as a density estimation. (b) Masked and unmasked performance of all methods measured by log KL divergence plus 1. Color blue represents masked datasets while yellow stands for unmasked datasets. The error bars show the variance. (c)-(h) UMAP embeddings of the following simulation methods with and without B cell subtype labels: Cytomulate, DG, LAMBDA, IG, scDesign2, and Splatter.

To gain further insights into the advantages of Cytomulate, we used UMAP for visual comparison among the different methods under the two settings. Although most methods except for Splatter yielded acceptable approximations to the real data when subtypes are given (Fig. 5), only Cytomulate was able to differentiate the underlying clusters with minimal distortion to their original shapes, even in the absence of detailed subtype information. Notably, LAMBDA suffered a severe loss of detail, approximating the entire dataset with a single Gaussian distribution centered between the two separated major clusters, where the actual density is low. Although scDesign2’s simulation came somewhat close to capturing the two major modes, noticeable distortion within each cluster remains. Given that KL divergence is a measure of distributional differences, our quantitative assessment, combined with the visual inspection, clearly shows Cytomulate’s advantage in effectively modeling heterogeneous data.

The robustness of Cytomulate makes it an appealing choice for practitioners, especially those who work with public datasets with existing cell types, since Cytomulate offers the greatest flexibility in terms of input. With reasonable cell types, Cytomulate can accurately capture the prominent characteristics of the data.

### Cytomulate is efficient

As a single CyTOF experiment could contain tens of thousands or millions of cell events along with 30 - 40 protein channels, computational efficiency including estimation and simulation is a significant issue. In fact, we initially included ZINB-WaVE (31), one of the top-performing scRNA-seq techniques in our comparison studies. However, it was excluded as it would take hours to simulate one moderately-sized dataset that contains around 50,000 cells. Specifically, we measured the estimation time (time required to construct the model and estimate the parameters) and simulation time (time needed to generate data from the estimated model) as our metrics for algorithm efficiency as all methods benchmarked involve these two steps. We conducted an analysis on the running time of each method by varying the number of cells and the number of markers using the CD8+ Naive T cells in the CyAnno dataset, which contains 8,414 cell events and 39 protein channels. To be more specific, while evaluating the impact of sample size, we used all 39 protein channels and applied bootstrap sampling to generate expression matrices containing various numbers of cell events. On the other hand, we fixed the number of cell events to 8,414 and sampled the columns to be included when investigating the effect of the number of markers on processing time. Besides including the six models previously described and benchmarked for accuracy, we also included Cytomulate with 5-component GMMs (Restricted Cytomulate) to gauge the impact of model selection on efficiency.

Overall, both estimation time and simulation time for all methods under consideration appear to be polynomial with respect to the number of cells and the number of protein channels (Fig. S3). Cytomulate is consistently faster than LAMBDA in all situations. For a moderate-size dataset containing 100,000 cells, even Cytomulate with model selection from 9 models takes around only 2 minutes (Fig. S3a). If restrictions are imposed, the estimation can be done within seconds. On the other hand, LAMBDA, due to the usage of a stochastic EM algorithm for the model estimation, is twice as slow, while scDesign2 is also among the slowest methods tested. When sample size reaches one million cells, both LAMBDA and scDesign2’s estimation times exceed one hour, which makes them not viable for large scale simulations. Surprisingly, LAMBDA and scDesign2 both suffer from poor scalability with regard to the number of markers as compared with other methods (Fig. S3b; Fig. S3d)

In terms of simulation time (Fig. S3c-d), all methods are relatively efficient as compared to estimation time. Despite the efficiency of all methods here, Cytomulate is always best with or without model selection, while Splatter is the least efficient. This characteristic allows Cytomulate to quickly generate many replicates using a single dataset as a reference, which is a common task. As a unique advantage to the Emulation mode, Cytomulate can perform parameter estimation using a small sample and then scale up by simulating a much larger sample. This further improves the scalability of Cytomulate given its efficiency in terms of simulation time. With the increasing capability of CyTOF to produce larger datasets and more channels, Cytomulate is fully capable of handling large datasets.

### Complex simulations

The core model of Cytomulate accounts for the expressions of each cell type as an essential part for simulation. Beyond the core model and unlike LAMBDA, Cytomulate can easily be extended to incorporate more complex effects commonly encountered in CyTOF experiments such as batch effects, temporal effects, as well as cell differentiations. While some of these effects are oftentimes considered nuisances with corrections desired, the ability to approximate them is crucial for methodological developments in the field of normalization or other pre-processing techniques. Also, the rising potential of trajectory inference with CyTOF datasets instead of scRNA-seq calls for a more thorough consideration of this topic. This section details the overall design and implementation of these complex simulation scenarios.

### Temporal effects

As discussed in (5), due to cell debris, etc. the sensitivity of a CyTOF machine may vary over time during data generation, causing signals for cells from one batch to fluctuate. In Cytomulate, we assume that the temporal effect for each batch is generated by a function defined on the unit interval such that the value at time zero is always zero, representing that there is no temporal effect at the beginning. We provide users with 3 options to simulate a temporal effect, each with a different level of control over the overall shape of the temporal effect:

1. Brownian Bridge (56): The shape of the entire trajectory is randomly generated from a Brownian Bridge where the ending value is randomly sampled from a normal distribution. To evaluate the function on the unit interval, the Akima spline (57) is used for interpolation. This option gives the users minimal control over the evolution of the temporal effects.
2. Polynomial: This option allows the user to specify the rough shape of the trajectory via the coefficients of a polynomial. A linear transformation will be used to make sure the resulting temporal effect starts at 0 and ends at a value randomly generated from a normal distribution.
3. Spline: During the preprocessing steps, it is possible that the users can extract the temporal effect component. The spline option permits the users to add the original temporal effect back to the simulated dataset via a smoothing spline. Since the size of the simulated dataset might differ from that of the original dataset, Cytomulate will automatically rescale the time interval so that the entire trajectory could be retained in the simulated dataset (Fig. 6a).

**Fig. 6.**
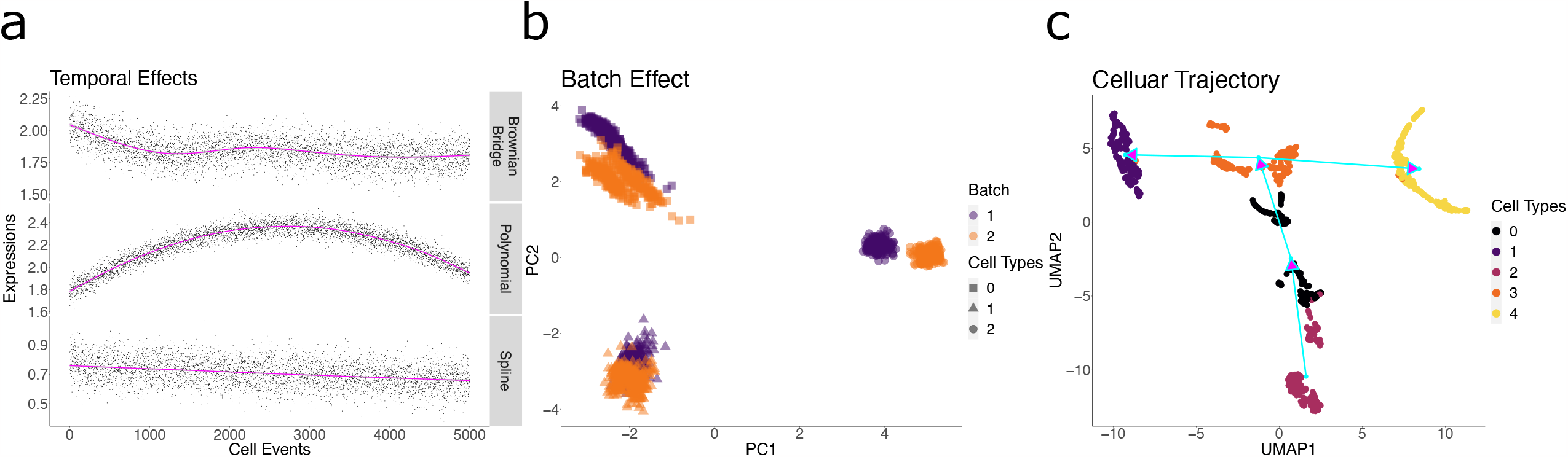
Examples of effects addressed in complex simulations. (a) Example temporal effects generated using the three methods: Brownian Bridge, Polynomial, and Spline. (b) Example of a simulated sample batch effect. Color difference represents batches, and point shape indicates cell types. (c) Example of the cell differentiation trajectory generated by Cytomulate. The trajectories are noted with light blue arrows.

### Batch effects

Previous research in the field (7), has indicated that the batch effects in CyTOF experiments vary not only by batches but also by cell types and protein channels. To simulate batch effects, we decompose the effects into global batch effects which assume different values for different batches and local batch effects which vary depending on the cell types and protein channels in a similar fashion to the way we decompose effects into main effects and interactions in Analysis of Variance (ANOVA). Further details can be found in the Materials and methods section. As a demonstration, we simulated two batches with 3 cell types using the creation mode of Cytomulate (Fig. 6b).

### Cellular trajectories

To simulate cellular trajectories, Cytomulate adopts a two-step procedure. In the first step, Cytomulate simulates the underlying graph of the cellular trajectories. Given the simulated graph, the actual protein expressions are then sampled in the second step.

Specifically, in the first step, a polytree (a directed acyclic graph whose underlying undirected graph is a tree) or a polyforest (a collection of nonoverlapping polytrees) is generated (58). A tree represents a single trajectory, where edges connect different cell types, allowing cells to differentiate along defined paths. In the Creation Mode, this is done by first randomly grouping the cell types and applying Kruskal’s algorithm (59) with random weights to sketch the underlying tree or forest. In the Emulation Mode, this is accomplished by the Clauset-Newman-Moore greedy modularity maximization algorithm (60) (61) followed by the Kruskal’s algorithm with the L2 distances among observed average cell means as weights. In both circumstances the root node represents the starting point of differentiation, where its child nodes inherit certain channels that define the entire lineage. To find the root node and convert the graph to a directed acyclic graph (DAG), a depth-first search with a randomly selected node as the root on each graph component is performed.

In the second step, Cytomulate starts the differentiation process. For selected channels that change between each pair of nodes, a Brownian Bridge combined with interpolating splines is used to generate the actual protein expressions along the polytree or the polyforest in a fashion similar to how Splatter (32) simulates paths. The time parameter of differentiation is determined by a Beta distribution. This scheme allows smooth transitions between cell types while also ensuring incremental changes along the tree (*i.e.* each tree contains a family of cell types with gradual changes). Further details can be found in the Materials and methods section. For illustration, we used the creation mode of Cytomulate to generate 5 cell types aligned on one polytree (Fig. 6c).

### Applications of Cytomulate within a unified framework

#### Framework overview

To ensure the interoperability of Cytomulate with downstream analyses and related workflows, we designed our software package using open standards in the field. Namely, our framework is expression matrix-driven with the optional support for PyCytoData—an easy-to-use interface for CyTOF and downstream analyses (Fig. 2b). Aside from two simulation modes, we support two main usage modes of Cytomulate. For users who prefer batch processing and working with servers, we recommend our command-line interface (CLI). Using a single command with the appropriate arguments and flags, users can generate their desired dataset and save it to disk in the format of their choice. Further, the CLI ensures scalability and interoperability by allowing users to utilize shell scripts and seamlessly integrate their results with other up- or downstream software. Alternatively, our interactive Python mode is designed for those who primarily work within Python and would benefit from the most flexibility. This interface supports fine-tuning of details while allowing users to explore the datasets interactively: from simulation to dimension reduction (DR) and plotting, a few lines of readable code are all they need.

With the wide adoption of CyTOF and the subsequent proliferation of analysis methods, a unified framework is increasingly important. While few methods, except ubiquitous tools such as PCA, have become virtually platform- and language-agnostic, many otherwise good software suffers from poor usability and extensibility. For example, in the process of evaluating Cytomulate, we found that LAMBDA falls far short of having a user-friendly interface: in fact, we spent a significant amount of time adapting the algorithm to our workflow for it to be viably assessed. To address these issues, we first developed the two usage modes as previously mentioned. The CLI uses the operating system’s shell as a common denominator to work with other parts of the workflow as necessary, whereas the interactive mode supports outputting a PyCytoData object. The latter brings the benefit of a suite of integrated downstream analysis tools and utility functions, including builtin normalization and CytofDR for DR evaluation (15). Beyond the software itself, our continuous integration and development workflows along with detailed documentation (https://cytomulate.readthedocs.io) ensure that Cytomulate is well maintained and easy for any user to adopt with minimal to no learning curve.

To demonstrate the usability and applicability of Cytomulate in CyTOF research, we hereby showcase three benchmark studies conducted using both Creation and Emulation modes. We also would like to emphasize the fact that Cytomulate’s potential goes much beyond the particular methods and benchmarks conducted in this section. With the flexibility of our framework and the open source initiative, other researchers can effortlessly extend the functionalities by simply using the PyCytoData object with their own workflows. A crucial advantage of employing Cytomulate instead of relying solely on real datasets is that users can specify characteristics of each sample, a task hardly achievable in real datasets without artificially subsampling and manipulations. Indeed, a proper benchmark requires the detailed controls necessary to distinguish between various effects, such as cell abundance and differentiation paths. Cytomulate not only supports all these configurations with both of our interfaces but also produces sensible results for practitioners as seen in the following benchmarks.

### Benchmarking DR using Cytomulate

As a practical application of Cytomulate, a previous DR benchmark study (15) included extensive usage of the creation mode to evaluate DR methods. Specifically, Wang et al. simulated 425 CyTOF datasets by specifying and systematically varying key features of CyTOF expressions. To further demonstrate the capabilities of Cytomulate for downstream analyses, we accessed and analyzed a subset of the results published on Database of Actionable Immunology’s (DBAI) (62–64) CytofDR Playground (https://dbai.biohpc.swmed.edu/cytof-dr-playground/). In particular, we chose samples with 100,000 cells and the following parameters: the number of markers {30, 35, 40}, the number of cell differentiation paths {2, 3, 5}, and the number of cell types {5, 10, 15, 20, 30}. These criteria yield 225 datasets in total with 25 replicates for each configuration. As for DR methods, we focused on the top-performing and most popular ones in the field according to the benchmarks: MDS (65), tSNE (16), UMAP (17), PCA (66), and SAUCIE (19).

Using two major categories (Global Structure Preservation and Local Structure Preservation) and the two subcategories of the Downstream Category, we validated the previous results that MDS is superior for structure preservation whereas UMAP and SAUCIE are better for clustering-based metrics (Fig. S4). These results are indeed concordant with results obtained from real datasets.Further, Wang et al. (15) demonstrated that Cytomulate produced good datasets for both downstream analyses and DR visualizations. While we defer to the original DR paper for those interested in the details of DR methods and their performances with regard to simulation settings, we would like to emphasize again the unified framework that enables this type of analysis. Users of Cytomulate can easily generate datasets and replicate the aforementioned DR results by using Cytomulate in conjunction with CytofDR through the PyCytoData framework.

### Validating clustering performance using Cytomulate

Clustering is a common and important workflow in the analysis of real CyTOF data because of its role in identifying similar cells. Many methods have been developed for CyTOF data, such as FlowSOM (11) and flowMeans (67). Traditional methods, such as K-Means (46), can in theory be applied to CyTOF data as well, but their performance is questionable. Two previous review papers (47,48) have extensively benchmarked popular clustering methods, and FlowSOM and flowMeans have in general been recommended. However, neither paper included simulation data as part of their workflows. To fill in the gap by incorporating Cytomulate as part of this analysis, we conducted a benchmark of three popular clustering methods to validate conclusions from the previous review papers.

In both Creation and Emulation modes, the ground truth labels are known, thus rendering Cytomulate a gold standard to assess whether clustering algorithms can correctly group the same cells in the same cluster. To demonstrate Cytomulate’s advantage, we simulated 20 datasets from the Levine_32dim dataset (40). Under the default setting with fixed probability of cell abundance (Fig. S5a), we found that flowMeans performs the best out of the three methods, achieving the highest Adjusted Rand Index (ARI) (68). FlowSOM follows closely behind flowMeans, whereas K-means lagged severely behind. To account for various cell abundances in real datasets, we also tested a setting by randomly varying the cell abundance probabilities generated from a Dirichlet distribution with parameters being a 14-dimensional vector of 0.1.

While the order of the three methods remained the same, K-means’s suboptimal characteristics for CyTOF data are exacerbated. Our results not only agree with the numerical analyses on Levine_32dim by Liu et al. (47), but they also highlight the usefulness of Cytomulate to simulate user-controlled characteristics of CyTOF, such as cell type abundances.

### Comparing batch normalization methods using Cytomulate

As part of Cytomulate’s complex simulation, users can easily simulate multiple batches of CyTOF datasets by adding batch effects. This procedure is designed to mimic the real situation in which practitioners have to account for systematic differences across multiple batches. With the development of many batch correction methods in the field, including ComBat (69), CytoNorm (7), and Harmony (70), it is important to assess how well each performs.

Cytomulate’s complex simulation provides a good basis for such a benchmark. We thus selected the aforementioned three methods and compared their performances using 10 paired simulated samples based on the Samusik (52) dataset. Within each sample, an “anchor” sample that is free of batch effect was first generated. A “replicate” sample was then generated with simulated batch effects.

To assess the extent to which batch effect is corrected, we computed the Earth Mover’s Distance (EMD) (71) between corrected batches, and smaller distance corresponds to a more effective batch normalization. Further, we created two scenarios: “High Variance” and “Low Variance” to represent two levels of potential amount of batch variation from sample to sample. In the case of “High Variance”, the batch effects were simulated from a normal distribution with zero mean and unit standard deviation. The standard deviation was set to 0.1 in the case of “Low Variance”. Across both high and low variance settings, CytoNorm is the best for CyTOF data, while Harmony is the worst. Also, the low variance setting yields overall smaller EMD as we expect (Fig. S5b). This benchmark shows again that methods designed specifically for CyTOF tend to outperform others, such as Harmony for scRNA-seq and ComBat for RNA-seq.

## Discussion and conclusions

Simulation is a powerful, flexible, and cost-effective tool for performance evaluation, which plays a vital role in methodological development. A common way to test analysis methods is to conduct simulation studies, where generating multiple datasets, with different parameters or assumptions, can be quickly accomplished with minimal cost. The lack of a formal method for accurately synthesizing CyTOF data warrants a dedicated statistical tool with careful considerations of CyTOF’s unique characteristics, such as its throughput and the correlation between protein channel expressions.

In this paper, we filled in the aforementioned void by introducing Cytomulate, a tool for simulating CyTOF data accurately and efficiently. It builds on Gaussian Mixture Models (GMM) with a fast and robust estimation procedure and appealing mathematical properties to approximate the real densities of CyTOF channels. Furthermore, Cytomulate features a well-defined probabilistic model to capture prominent aspects of CyTOF data while also incorporating two simulation modes: creation mode and emulation mode to cater to the needs of various researchers. By thoroughly benchmarking Cytomulate’s estimation and simulation efficiency, resemblance to real data, and robustness again prior cell-type resolution on six publicly available datasets, we demonstrated that Cytomulate is superior in approximating the overall data distributions regardless of the granularity of the input. Our algorithm further has the versatility of simulating more complicated phenomena commonly encountered in a CyTOF experiment such as signal drifts (temporal effects), batch effects, and cellular trajectories, which are partially or totally unsupported by other tools. Finally, we showcased Cytomulate’s integration with PyCytoData as an open standard for CyTOF data processing in Python while also highlighting Cytomulate’s usage in a series of benchmark studies involving several DR methods (15), clustering algorithms, and batch correction tools. By integrating the philosophy of user-centric software and the superior performance of our probabilistic model, Cytomulate offers a practical step towards addressing the usability problem in the field (15,45).

Potential limitations of this study involve the estimation procedure we chose for the emulation mode. Since the parameters are estimated for each cell type separately, in the presence of high-dimensional data, the usage of the EM algorithm could render the performance of a GMM unsatisfactory when the number of cell events is low. A potential remedy for rare cell types would be to borrow “strength” from other cell types to stabilize the parameter estimation via a Bayesian hierarchical setup. What’s more, as a trade-off for efficiency, various aspects of CyTOF data such as means, covariances, and cell differentiations are captured in a sequential manner instead of being modeled jointly. Consequently, although the GMMs provide a reasonable approximation to the data distribution of CyTOF due to its universal approximation to densities (41), the estimated parameters are likely biased. Due to the sheer number of cells typically encountered in CyTOF and the complexity of the model, traditional inference methods such as Markov Chain Monte Carlo (MCMC) might not ameliorate this situation. Fortunately, tremendous progress has been made in the field of deep generative models, which has been successfully applied to tasks such as estimating densities and synthesizing realistic-looking images on large datasets (72,73). Some typical examples include normalizing flow (74), variational auto-encoder (75), and generative adversarial networks (76). Exploiting said methods to gain deeper insights into CyTOF datasets thus could be a possible future direction.

Another future direction constitutes the modeling of clinical phenotypes based on simulated CyTOF datasets. Namely, suppose we denote *y* as the phenotype and *x* as CyTOF data, and an interesting direction is to simulate *y* from *p*(*y*|*x*). Currently, Cytomulate has the ability to generate CyTOF datasets from *p*(*x*). Adding the connection between simulated datasets and phenotypes will increase the applicability and practicality of Cytomulate in real settings. We are optimistic that our simulation framework as a first in the field will spark further interests in these exciting directions, and our model will serve as a reference for future works.

## Materials and methods

### Probabilistic model of Cytomulate

The probabilistic model of Cytomulate is intended to model the *arcsinh*(./5)-transformed CyTOF data. In this section, we will gradually build up towards the final model aided by what we observed in the section on characteristics of CyTOF data.

### Basic model

We will start with the simplest scenario where we only have one batch, *n* = 1, ···, *N* cell events on only one cell type with measurements on *m* = 1, ⋯, *M* protein channels.

We start with a continuous latent variable for the protein expressions:

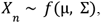

where *f*(μ, Σ) denotes some distribution with an *M*-by-1 matrix of mean μ, and an *M*-by-*M* matrix of variance-covariance matrix Σ. Given the near-Gaussianity nature of CyTOF data, we use a Gaussian Mixture Model to approximate this distribution in Cytomulate.

Since protein expressions should be non-negative, we further truncate the variable below zero. Mathematically, this can be accomplished by

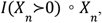

where ° is the Hadamard product, *I* is an indicator function, and ≻ is a component-wise inequality.

Finally, we incorporate the machine noise into the final protein expressions resulting in the basic model of Cytomulate:

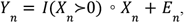

where *E*_*n*_ ∼ *L* and *L* is some error distribution.

### Core model

The core model of Cytomulate is only a slight generalization upon the basic model by considering *p* = 1, ⋯, *P* cell types in one batch.

We denote the probability of observing the *p*th cell type by π_*p*_. Then, for the *n*th cell event, we assume that the cell type is sampled from

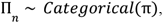

where π = [π_1_, ···, π _*P*_] is a *P*-by-1 probability vector.

Suppose the cell type for the *n*th cell event is *p*.We then update the notation for the latent variables accordingly:

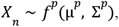

where *f*(μ^p^, Σ ^*p*^) denotes some distribution with mean μ^*p*^, and variance-covariance matrix Σ^*p*^.

The computation of the final protein expressions remain unchanged:

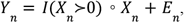

where *E _n_* ∼ *L* and *L* is some error distribution.

### Complex simulations

#### Batch effects

To add batch effects, suppose we have *b* = 1, ···, *B* batches, *m* = 1, ⋯, *M* protein channels, and *p* = 1, ⋯, *P* cell types. For the *b*th batch, we have *n* = 1, ···, *N*^b^ cell events.

Similar to the setup in the core model, we assume that the cell type of the *n*th cell event in the *b*th batch is sampled from

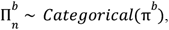

where 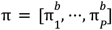 is a *P*-by-1 probability vector. The corresponding latent variable is then denoted by 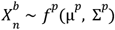. Note that while we allow the cell abundances to vary among batches, we only use one distribution for one cell type.

With this setup, we introduce a set of random vectors, each representing the batch effect on the *p* th cell type in the *b*th batch:

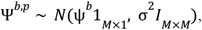

where

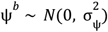

represents the center around which the batch effects on the cell types and protein channels in the

*b*th batch vary. We further add two sets of constraints to ensure the identifiability of the parameters:

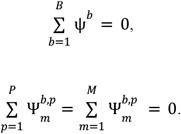

#### Temporal effects

Suppose we have *b* = 1, ···, *B* batches, and *m* = 1, ⋯, *M* protein channels. For the *b*th batch, we have *n* = 1, ···, *N*^*b*^ cell events. To add temporal effects to the *n*th cell event in the *b*th batch, we assume that there is a stochastic process:

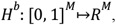

such that *H*^*b*^ (0) = 0 This reflects our belief that there should be no temporal effects at the beginning of an experiment. Denote the realization of *H*^*b*^ by Φ^*b*^, then the temporal effect on the *n*th cell event is calculated as:

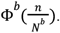

#### Cellular trajectories

The setup of simulating cellular trajectories is a little more involved. Again, we assume that we have *b* = 1, ···, *B* batches, *m* = 1, ⋯, *M* protein channels, and *p* = 1, ⋯, *P* cell types. For the *b* th batch, we have *n* = 1, ···, *N*^*b*^ cell events.

To get started, we first fix cell type *p*. We suppose that it differentiates into *z*^*p*^ “children” denoted by

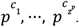

In Cytomulate, we ensure that the collection of “parent-children” relationships form a polytree (a directed acyclic graph whose underlying undirected graph is a tree) or a polyforest (a collection of nonoverlapping polytrees) (58).

To accomplish this, a weighted complete undirected graph with cell types as nodes are constructed. In the Creation Mode, the weights on the edges are randomly assigned. In the Emulation Mode, the L2 distances among observed average protein expressions of all cell types are used as weights.

Next, we carve out a spanning tree or a spanning forest via the Clauset-Newman-Moore greedy modularity maximization algorithm (60) (61) followed by the Kruskal’s algorithm (59). Finally, to convert the graph to a directed acyclic graph (DAG), a depth-first search with a randomly selected node as the root on each graph component is performed.

Now fix the “child” cell type *p*^*c*^. We construct the corresponding cellular trajectory from cell type *p* to cell type *p*^*c*^ in a similar fashion to how we construct temporal effects. We start with a stochastic process

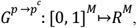

such that the *m*th component satisfies

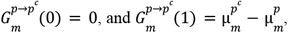

where 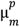 and 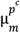 are the mean expression levels of the *m*th protein channel of cell type *p* and cell type *p*^*c*^, respectively. Given these two constraints, a Brownian Bridge becomes an intuitive choice for the task of trajectory generation since it is a Brownian Motion with an extra condition on where the position of the process should be at time 1. To simulate such a stochastic process that starts at 0 and ends at 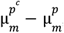, we utilize a standard Brownian Motion *B*_*t*_ as the basic building block and simply let 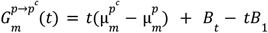. This stochastic process is also adopted by Splatter (32) in generating the actual realizations of cellular trajectories denoted here by 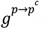. When generating the final protein expressions for the *n*th cell event in the *b*th batch whose cell type is *p*, we first sample the “child” cell type *p*^*c*^ from

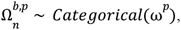

where 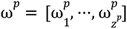 by-1 probability vector. Notice that the parameters of the distribution only vary with cell types.

We then sample its *m*th component of the actual position 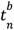 on the cellular trajectory 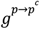

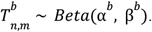

### Final model

With these setups, the final protein expressions for the *n*th cell event in the *b*th batch whose cell type is *p* and who is differentiating into cell type *p*^*c*^ can be simulated via the following two steps:

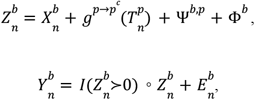

where 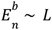 and *L* is some error distribution.

### Estimation procedure for the emulation mode

Given the expression matrix along with the cell type label for each cell event, Cytomulate first groups the observations by cell types and associates with each type a GMM whose parameters are then estimated via the EM algorithm adopted by the *scikit-learn* Python package (77). To further improve the approximation of a GMM to CyTOF data, for each cell type, we match the probability of obtaining a near-zero value estimated by the model with the actual frequency observed in the data. This is done by randomly zeroing out simulated values with probabilities inversely proportional to their magnitudes.

### Evaluation metrics

In this section, we elaborate on the metrics we used in the comparison studies. Suppose in the simulated batch we have *p* = 1, ⋯, *P* cell types. Then, for the *p*th cell type, we calculate the sample mean *m*_*p*_ and the sample covariance as *c*_*p*_. We carry out the same procedure on the observed dataset to the corresponding 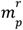 and 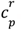. The L2 distance between the simulated and observed means is found via 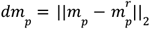. The Frobenius distance (78) between the simulated and observed covariances is calculated as 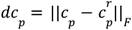. The KL divergence *dk*_*p*_ is estimated using the method proposed in (54); the pMSE *pmse*_*p*_ is calculated using the R package *synthpop* (55). Finally, the discrepancy measures for all the cell types are aggregated using a weighted average where the weights are determined by the cell abundances.

### Software implementation details

All analyses, plotting, and benchmarks were performed using the R (4.0.2) and Python (3.8.8) programming environments in MacOS 12.4. All default settings were used for Cytomulate, Splatter, and scDesign2. For LAMBDA, we skipped the multinomial sampling process given that all cell types were known.

Cytomulate is implemented and supported in Python (v3.5 or later), and the CLI is available on all platforms and systems where a compatible python interpreter is present with the proper package installations. The detailed documentation and tutorials are freely available and hosted on the cloud using ReadTheDocs (https://cytomulate.readthedocs.io). All software features have been thoroughly tested with the *pytest* framework.

For the efficiency benchmarks, we carried out the experiment on a MacBook Pro with a 2.9 GHz Quad-Core Intel Core i7 processor running MacOS 13.0.1. Since Cytomulate is implemented in Python while Splatter, LAMBDA, and scDesign2 are R packages, we used Python’s time module to measure the performance of Cytomulate-related methods and Sys.time function in R to time Splatter, LAMBDA, and scDesign2. All times are reported in seconds.

### Statistical analyses

We employed Pearson correlation coefficients for all correlation matrices and plots. For all order-based metrics, higher orders represent better performance. In cases where ties occurred, we used the average order of tied values. For all boxplots appearing in this study, box boundaries represent interquartile ranges, whiskers extend to the most extreme data point which is no more than 1.5 times the interquartile range, and the line in the middle of the box represents the median. All barplots showing discrepancies among all methods under comparison on various evaluation metrics were plotted on the logarithm scale. To increase visual clarity, we added one to all the data points before plotting so that the results would all be positive. Error bars with width being two standard deviations were also added to further facilitate comparison.

## Declarations

### Ethics approval and consent to participate

Not applicable

### Consent for publication

Not applicable

### Availability of data and materials

The Levine_32dim, Levine_13dim (38), and Samusik (52) datasets can be accessed from the HDCytoData package in R (https://github.com/lmweber/HDCytoData) (40). The CyAnno (FlowRepository ID: FR-FCM-Z2V9) (39,79) and LG (GEO accession: GSE99093; FlowRepository ID: FR-FCM-ZY4P) (53,79,80) can be accessed from their respective publications. The Covid cohort is hosted on DBAI (https://dbai.biohpc.swmed.edu/) (62–64) with the following accession codes: CyTOF_00000000000001 to CyTOF_00000000000012.

The Cytomulate algorithm is available at cytomulate.readthedocs.io, and also shared through the Database of Actionable Immunology (dbai.biohpc.swmed.edu) (62–64). All the datasets as well as the source code used to generate the findings are available in Zenodo (https://doi.org/10.5281/zenodo.10005508) and are released under the MIT license (81).

### Authors’ contributions

Y.Y. performed all analyses. Y.Y. and K.W. created the *Cytomulate* Package. Z.L provided analysis pipelines for LAMBDA. X.W. conceived the study. T.W. and X.W. designed and supervised the whole study. All authors wrote the manuscript.

## Acknowledgements

Not applicable

## Supplementary information

### Additional file 1

**Table S1:** A table that provides the dataset name, species, the anatomic site, and its source for accession.

### Additional file 2

**Fig. S1 -** Examples of complex characteristics observed in real datasets.

**Fig. S2 -** Detailed results on embeddings of the original and simulation datasets.

**Fig. S3 -** Runtime benchmarks of all simulation methods in the study.

**Fig. S4 -** Performance of DR methods benchmarked with Cytomulate’s creation mode.

**Fig. S5 -** Clustering and batch correction benchmark results using Cytomulate.

