## Additional file1 for "Cytomulate: accurate and efficient simulation of CyTOF data"

| Dataset | Species | Anatomic Site | Source | Number of Markers | Number of Cells Events | Number of Cell Types |
|---|---|---|---|---|---|---|
| Levine_32dim | Human | Bone Marrow | HDCytoData | 32 | 72,463 | 32 |
| Levine_13dim | Human | Bone Marrow | HDCytoData | 13 | 167,044 | 24 |
| Samusik | Mouse | Bone Marrow | HDCytoData | 39 | 53,173 | 24 |
| CyAnno | Human | Peripheral Blood | https://flowrepository.org/id/FR-FCM-Z2V9 | 39 | 123,033 | 39 |
| Covid | Human | Peripheral Blood | https://dbai.biohpc.swmed.edu/cytof-database.php | 43 | 208,712 | 21 |
| LG | Mouse | Lacrimal Gland | http://flowrepository.org/id/FR-FCM-ZY4P | 31 | 284,055 | 7 |

**Table S1.** A table that provides the dataset name, species, the anatomic site, and its source for accession.
