## Additional file 2 for "Cytomulate: accurate and efficient simulation of CyTOF data"

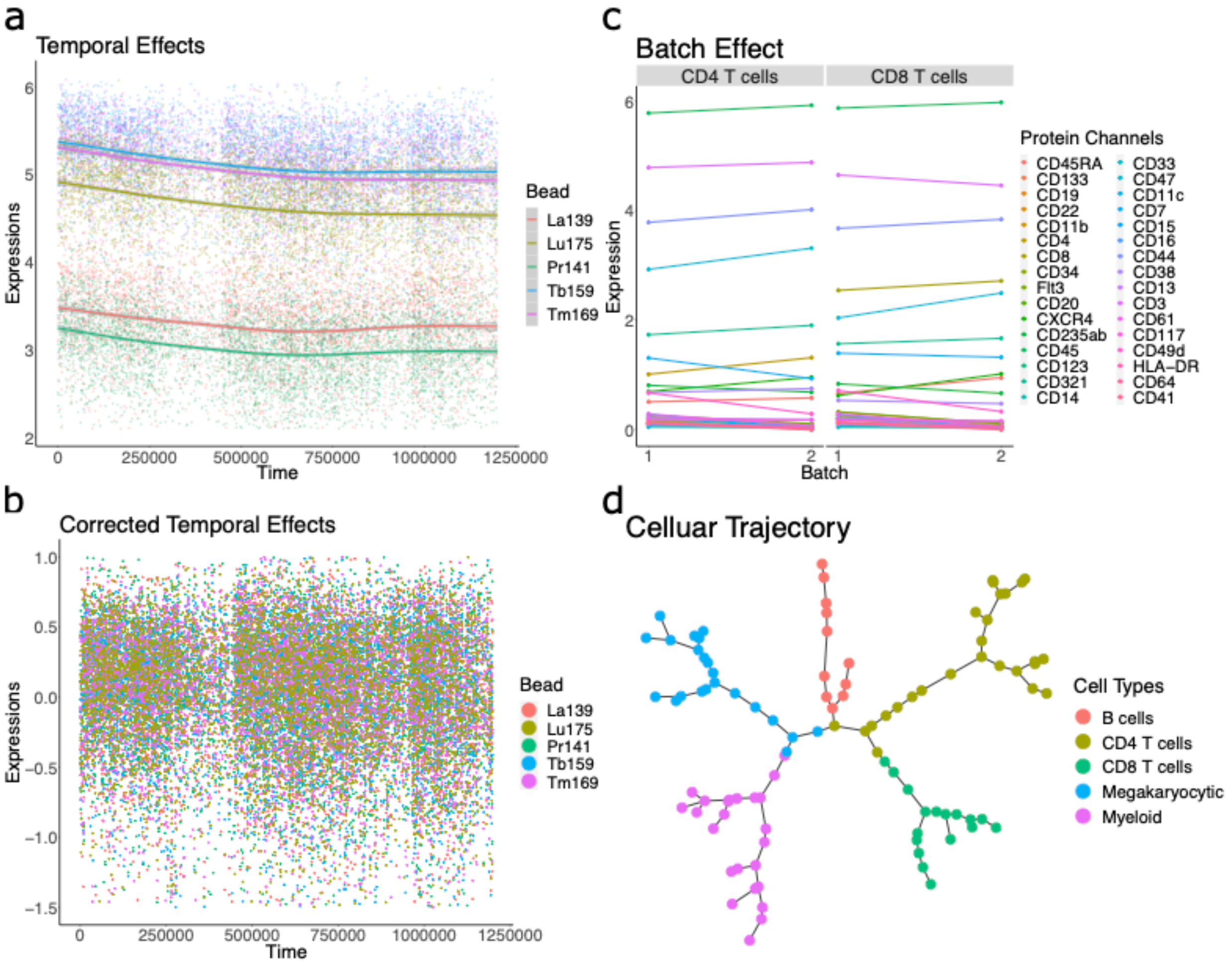


**Fig. S1.** Examples of complex characteristics observed in real datasets. (a) The temporal effect present in the real PBMC Finck dataset, as measured by the signal drift of five bead channels. Colors represent the different bead channels. (b) The corrected time effects after bead normalization. Colors represent the different bead channels. (c) The batch effect of the two patients in Levine_13dim visualized with an interaction plot in which the mean expression of protein channels change from sample to sample. Each color is a protein marker. (d) An example cellular trajectory fitted using the CytoTree package based on a real Bone Marrow dataset from the CytoTree documentation. Each node is a meta-cell colored by the cell type, and each edge represents the path.

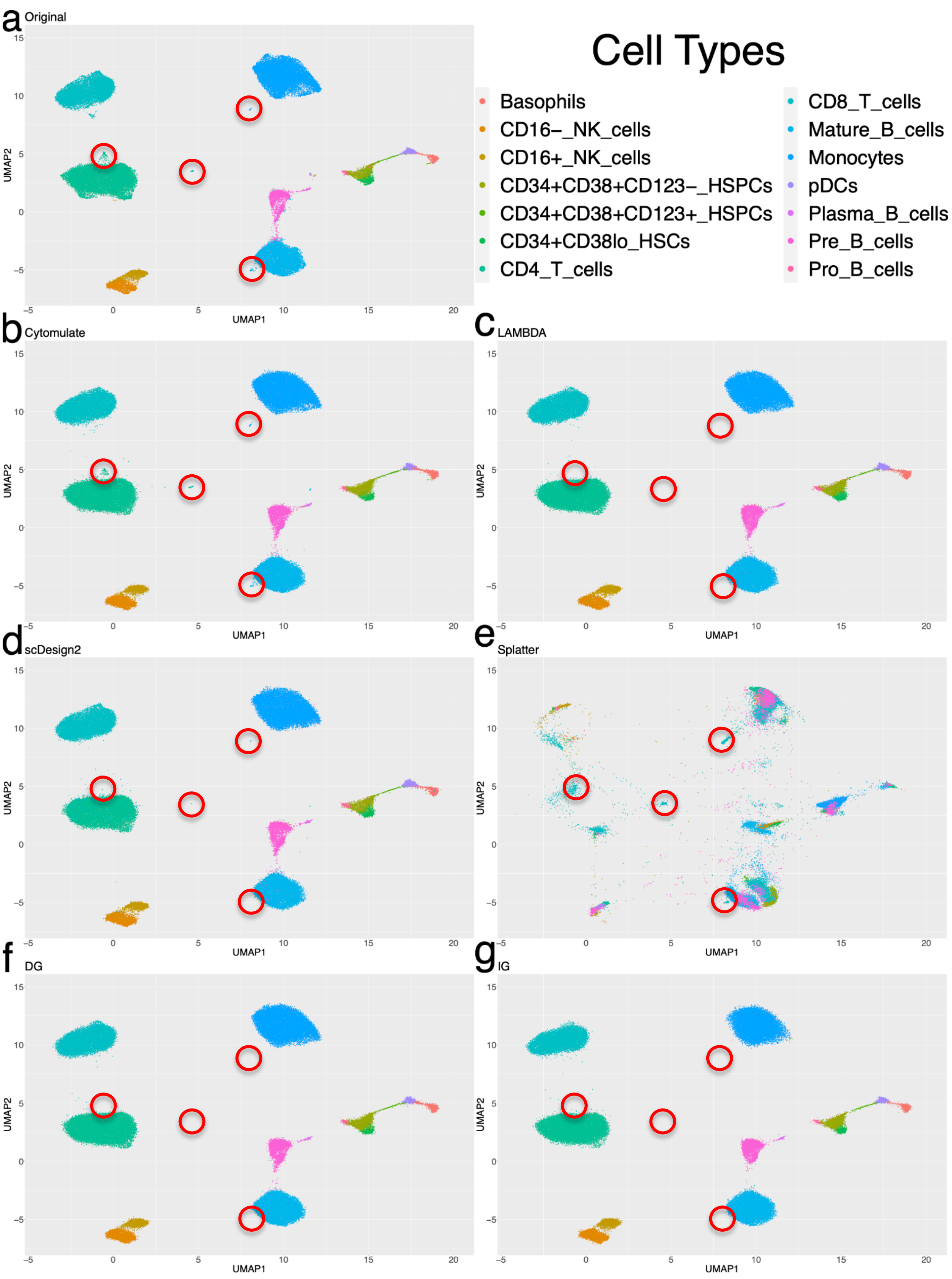

a Original
b Cytomulate
c LAMBDA
d scDesign2
e Splatter
f DG
g IG
UMAP1
UMAP2
Cell Types
Basophils
CD16−_NK_cells
CD16+_NK_cells
CD34+CD38+CD123−_HSPCs
CD34+CD38+CD123+_HSPCs
CD34+CD38lo_HSCs
CD4_T_cells
CD8_T_cells
Mature_B_cells
Monocytes
pDCs
Plasma_B_cells
Pre_B_cells
Pro_B_cells

**Fig. S2.** Detailed results on embeddings of the original and simulation datasets. (a) The UMAP embedding of the original Levine_32dim dataset. In (b), (c), (d), (e), (f), and (g), the UMAP embeddings of simulated datasets generated by Cytomulate, LAMBDA, scDesign2, Splatter, DG, and IG are plotted respectively. Each point represents a cell in the embedding space, and cells are colored by their cell types. Red circles are overlaid to pinpoint the locations of small clusters present in the original dataset.

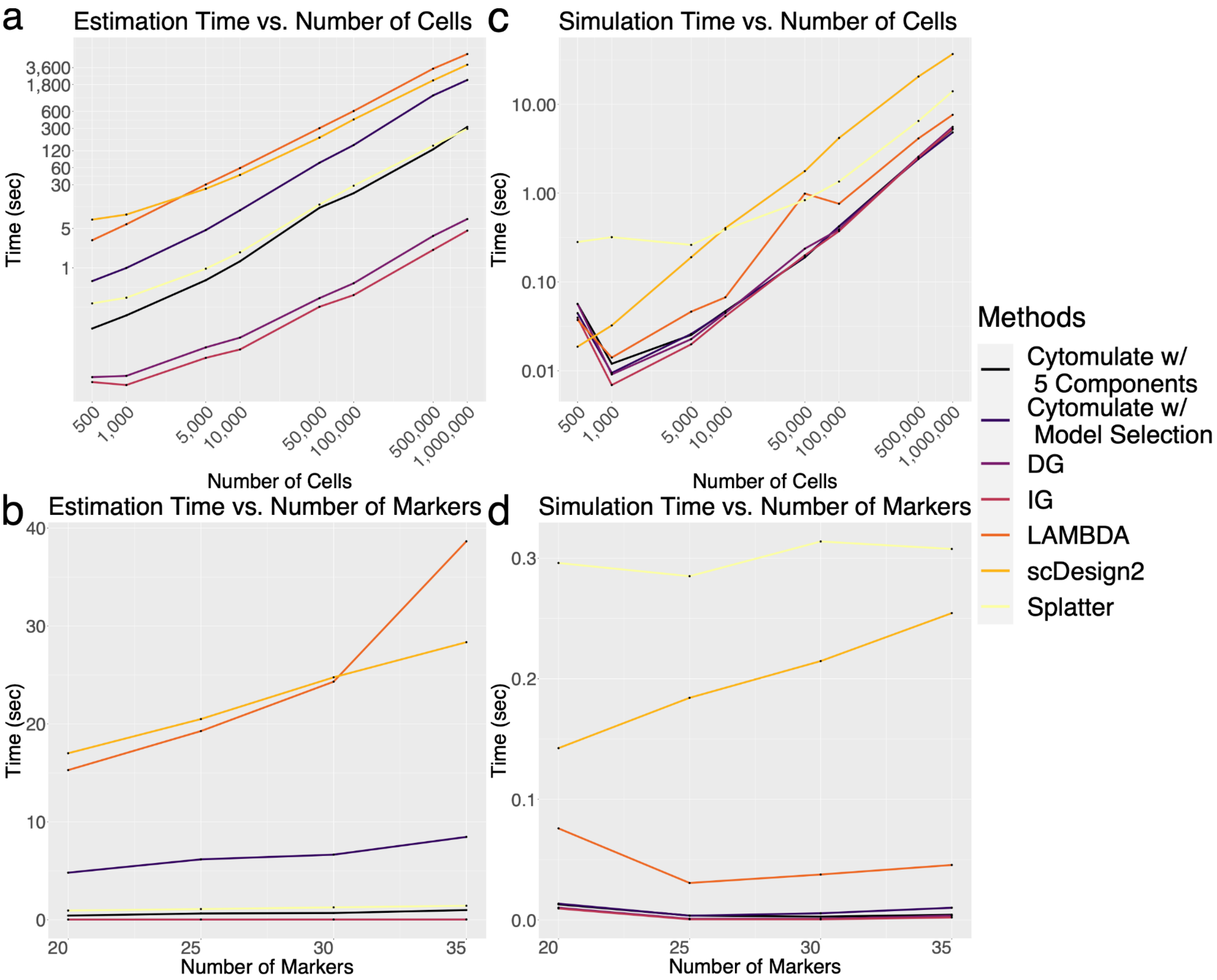


**Fig. S3.** Runtime benchmarks of all simulation methods in the study. In (a) and (b), we measured the estimation time of each method with various numbers of cells and markers respectively. The estimation time includes construction of the model object and estimation of parameters used in each model based on the dataset given. In (c) and (d), we benchmarked the simulation time of each method against different numbers of cells and markers. Simulation time is defined as the time necessary to generate data from the estimated parameters.

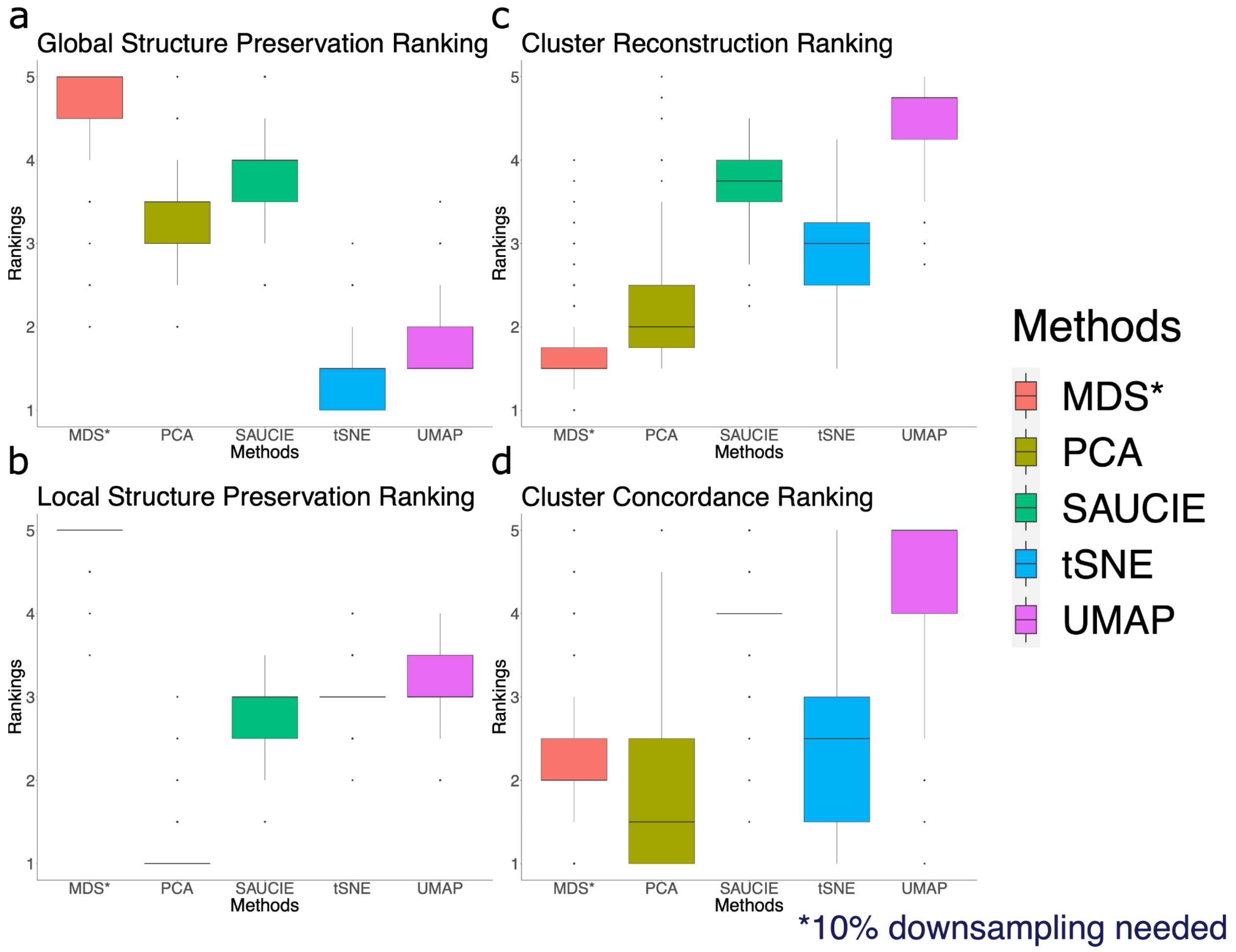


**Fig. S4.** Performance of DR methods benchmarked with Cytomulate's creation mode. Five methods are included, and all rankings are accessed from CytofDR Playground. "*" indicates 10% downsampling of the datasets used for DR and evaluation. (a) Rankings of DR methods using global structure preservation metrics. (b) Rankings of DR methods using local structure preservation metrics. (c) Rankings of DR methods using cluster reconstruction metrics. (d) Rankings of DR methods using cluster concordance metrics.

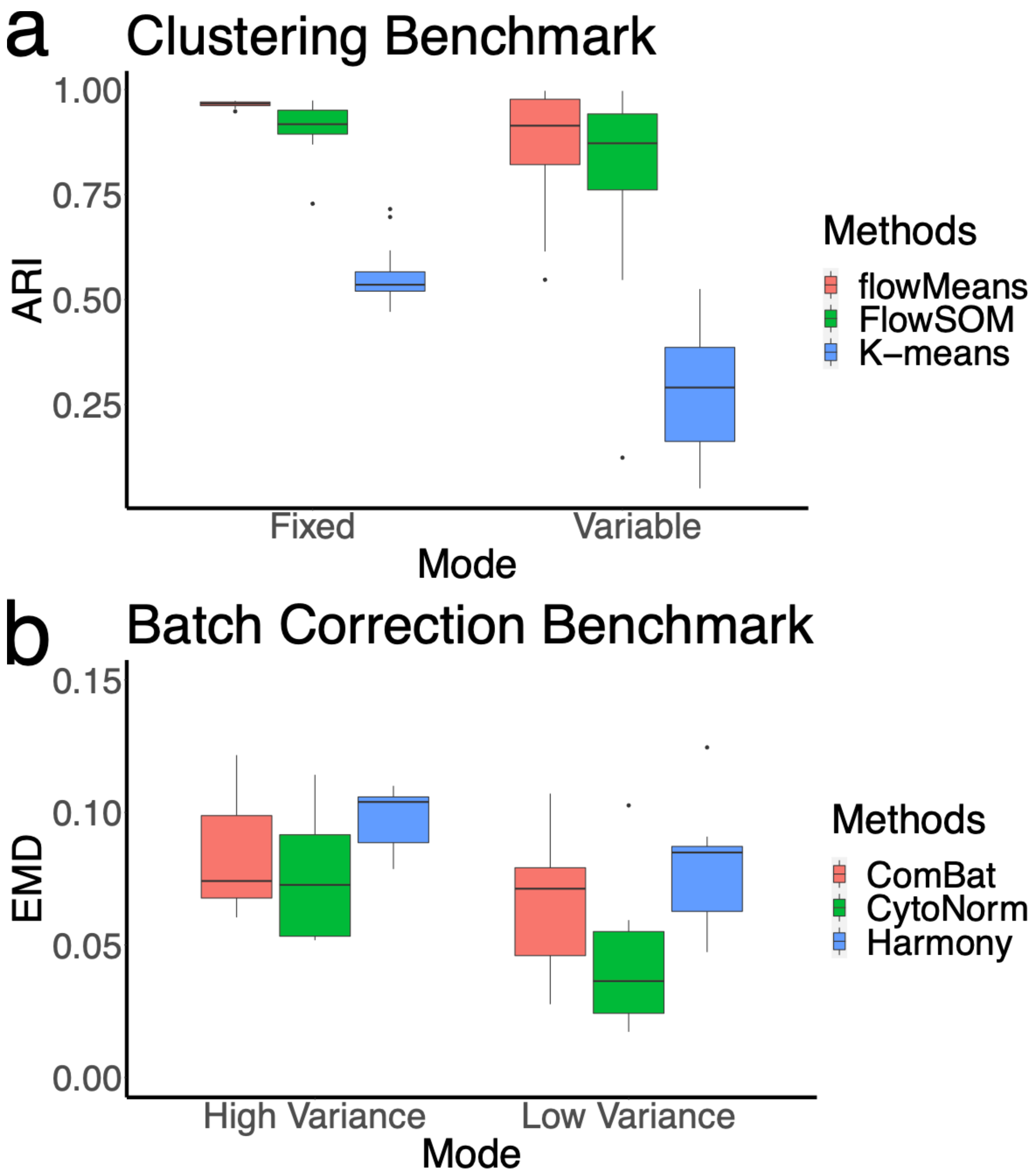


**Fig. S5.** Clustering and batch correction benchmark results using Cytomulate. (a) The Adjusted Rand Index (ARI) of three clustering methods across 20 replicates. Higher is better. (b) The Earth Mover's Distance (EMD) between batches for three batch correction methods benchmarked across 10 replicates. Smaller is better.
